## Supplemental Figures S1-S18 for "A plant-like mechanism coupling m6A reading to polyadenylation safeguards transcriptome integrity and developmental genes partitioning in *Toxoplasma*"

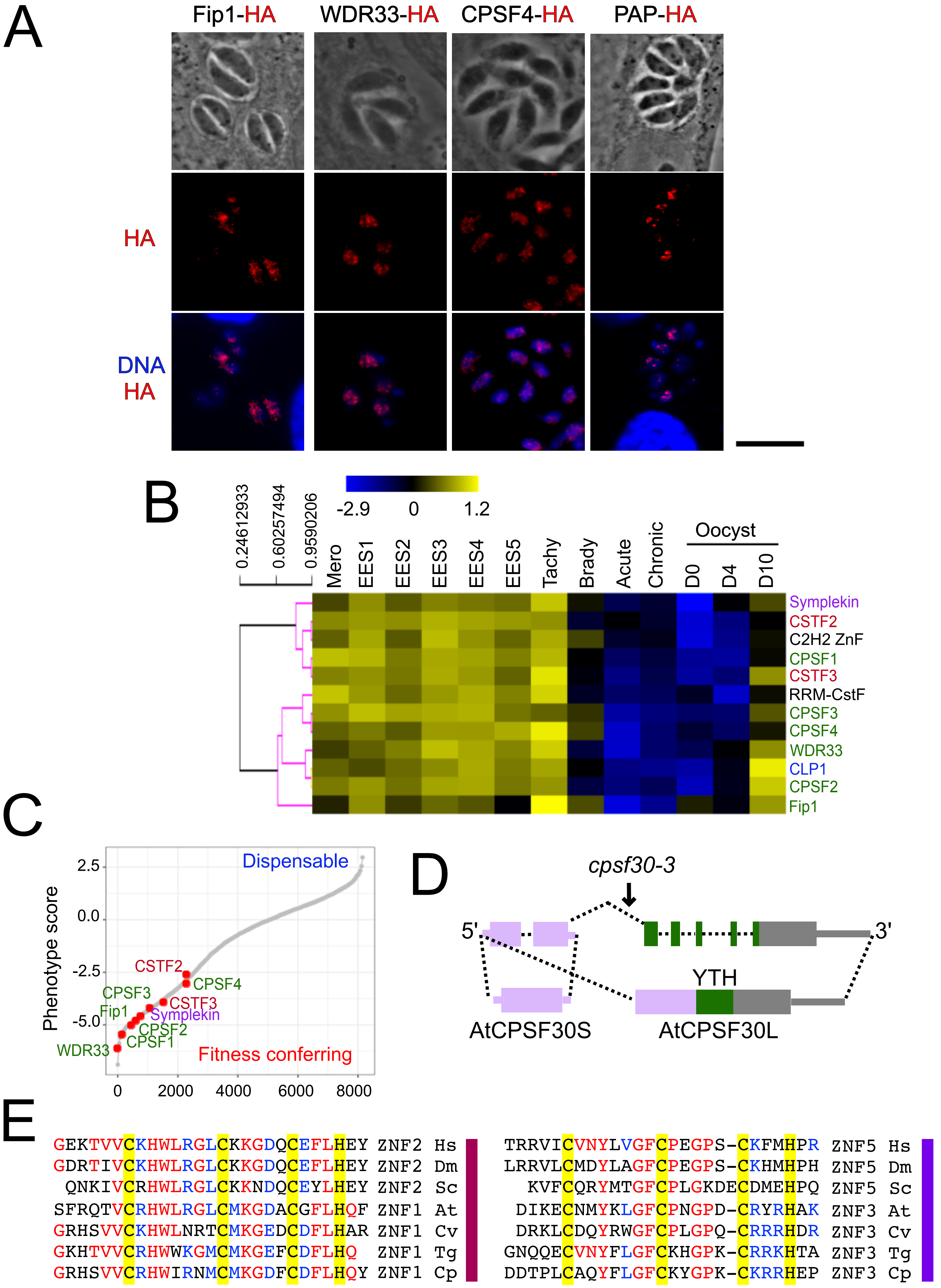
A
Fip1-HA
WDR33-HA
CPSF4-HA
PAP-HA
HA
DNA
HA
B
0.24612933
0.60257494
0.9590206
-2.9
0
1.2
Mero
EES1
EES2
EES3
EES4
EES5
Tachy
Brady
Acute
Chronic
Oocyst
D0
D4
D10
Symplekin
CSTF2
C2H2 ZnF
CPSF1
CSTF3
RRM-CstF
CPSF3
CPSF4
WDR33
CLP1
CPSF2
Fip1
C
Phenotype score
2.5
0.0
-2.5
-5.0
0
2000
4000
6000
8000
Dispensable
Fitness conferring
CSTF2
CPSF4
CPSF3
CSTF3
Fip1
Symplekin
CPSF2
CPSF1
WDR33
D
cpsf30-3
5'
3'
YTH
AtCPSF30S
AtCPSF30L
E
GEKTVVCKHWLRGLCKKGDQCEFLHEY ZNF2 Hs
GDRTIVCKHWLRGLCKKGDQCEFLHEY ZNF2 Dm
QNKIVCRHWLRGLCKKNDQCEYLHEY ZNF2 Sc
SFRQTVCRHWLRGLCMKGDACGFLHQF ZNF1 At
GRHSVVCKHWLNRTCMKGEDCDFLHAR ZNF1 Cv
GKHTVVCRHWWKGMCMKGEFCDFLHQ ZNF1 Tg
GRHSVVCRHWIRNMCMKGDFCDFLHQY ZNF1 Cp
TRRVICVNYLVGFCPEGPS-CKFMHPR ZNF5 Hs
LRRVLCMDYLAGFCPEGPS-CKHMHPH ZNF5 Dm
KVFCQRYMTGFCPLGKDECDMEHPQ ZNF5 Sc
DIKECNMYKLGFCPNGPD-CRYRHAK ZNF3 At
DRKLCDQYRWGFCPLGPQ-CRRRHDR ZNF3 Cv
GNQQECVNYFLGFCKHGPK-CRRKHTA ZNF3 Tg
DDTPLCAQYFLGFCKYGPK-CKRRHEP ZNF3 Cp

**Sup. Fig. 1**

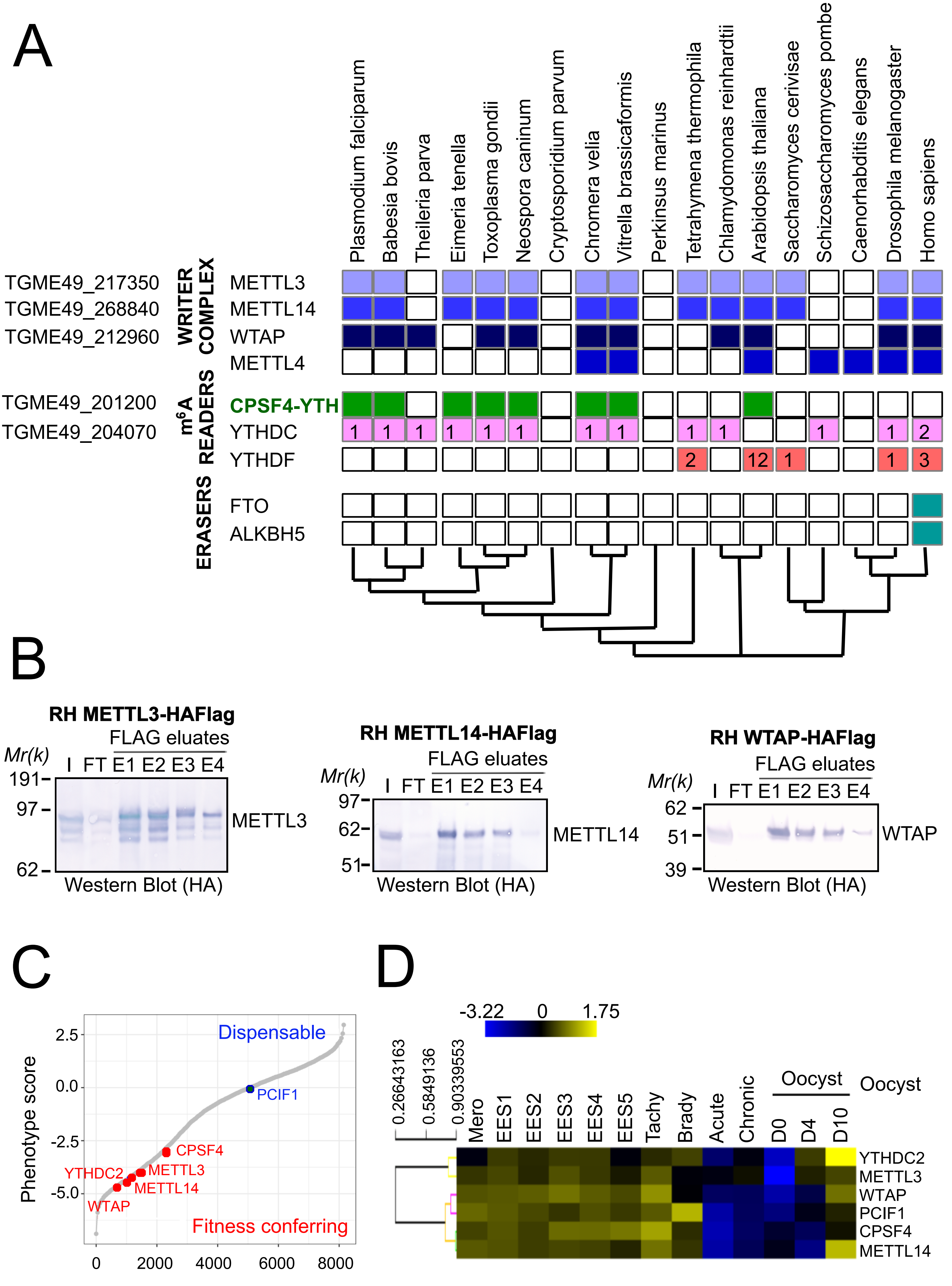

A
Plasmodium falciparum
Babesia bovis
Theileria parva
Eimeria tenella
Toxoplasma gondii
Neospora caninum
Cryptosporidium parvum
Chromera velia
Vitrella brassicaformis
Perkinsus marinus
Tetrahymena thermophila
Chlamydomonas reinhardtii
Arabidopsis thaliana
Saccharomyces cerivisae
Schizosaccharomyces pombe
Caenorhabditis elegans
Drosophila melanogaster
Homo sapiens
WRITER COMPLEX
TGME49_217350 METTL3
TGME49_268840 METTL14
TGME49_212960 WTAP
METTL4
m6A READERS
TGME49_201200 CPSF4-YTH
TGME49_204070 YTHDC
YTHDF
ERASERS
FTO
ALKBH5
B
RH METTL3-HAFlag
FLAG eluates
Mr(k)
I FT E1 E2 E3 E4
191
97
62
METTL3
Western Blot (HA)
RH METTL14-HAFlag
FLAG eluates
Mr(k)
I FT E1 E2 E3 E4
97
62
51
METTL14
Western Blot (HA)
RH WTAP-HAFlag
FLAG eluates
Mr(k)
I FT E1 E2 E3 E4
62
51
39
WTAP
Western Blot (HA)
C
Phenotype score
Dispensable
PCIF1
CPSF4
YTHDC2
METTL3
METTL14
WTAP
Fitness conferring
2.5
0.0
-2.5
-5.0
0
2000
4000
6000
8000
D
-3.22
0
1.75
0.26643163
0.5849136
0.90339553
Mero
EES1
EES2
EES3
EES4
EES5
Tachy
Brady
Acute
Chronic
Oocyst
D0
D4
D10
Oocyst
YTHDC2
METTL3
WTAP
PCIF1
CPSF4
METTL14


**Sup. Fig. 2**

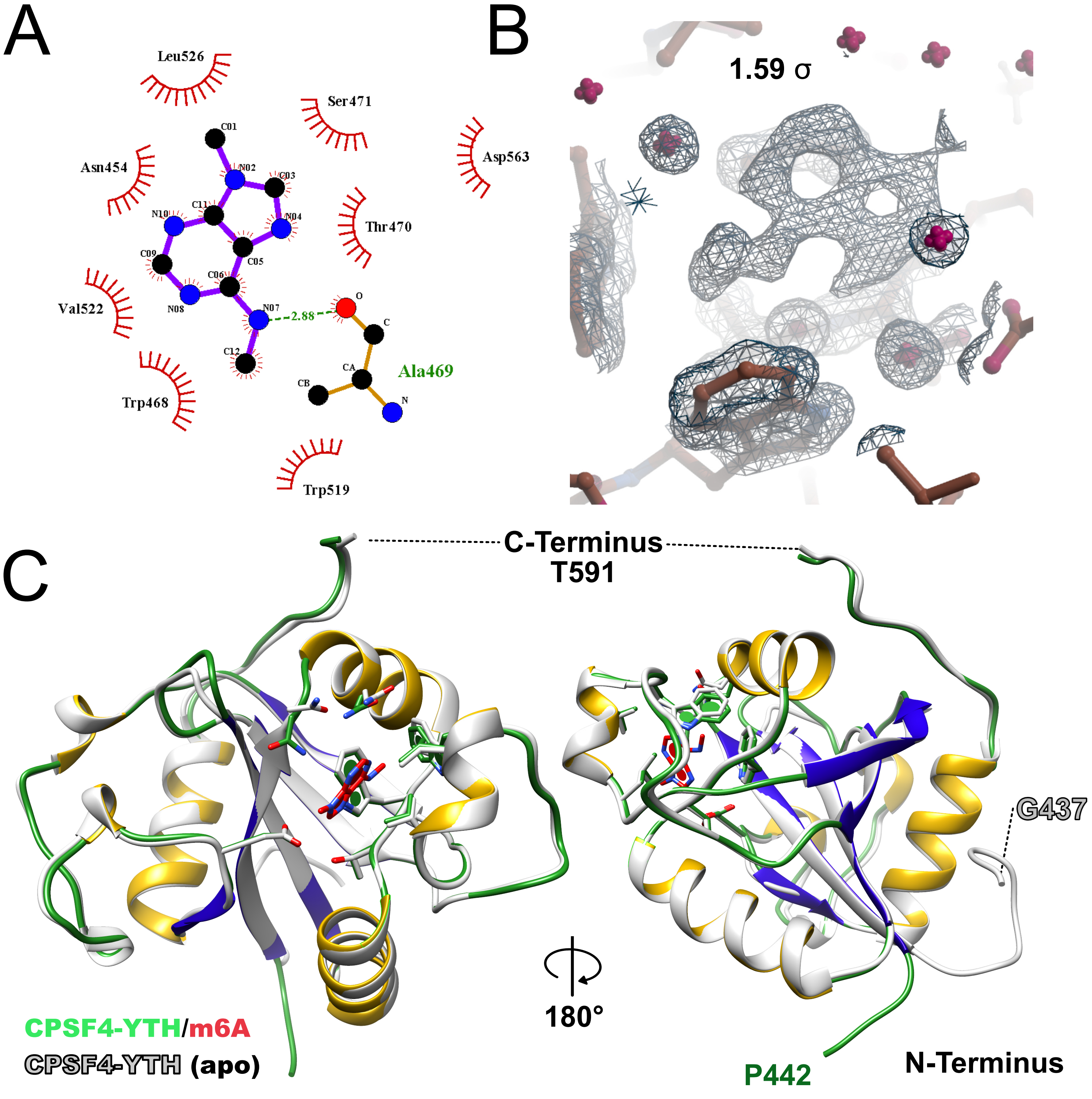
A
Leu526
Ser471
Asp563
Asn454
Thr470
Val522
Trp468
Trp519
Ala469
2.88
C01
N02
C03
N04
C05
C06
N07
N08
C09
N10
C11
C12
O
C
CA
CB
N
B
1.59 σ
C
C-Terminus
T591
G437
180°
CPSF4-YTH/m6A
CPSF4-YTH (apo)
P442
N-Terminus

### Sup. Fig. 3

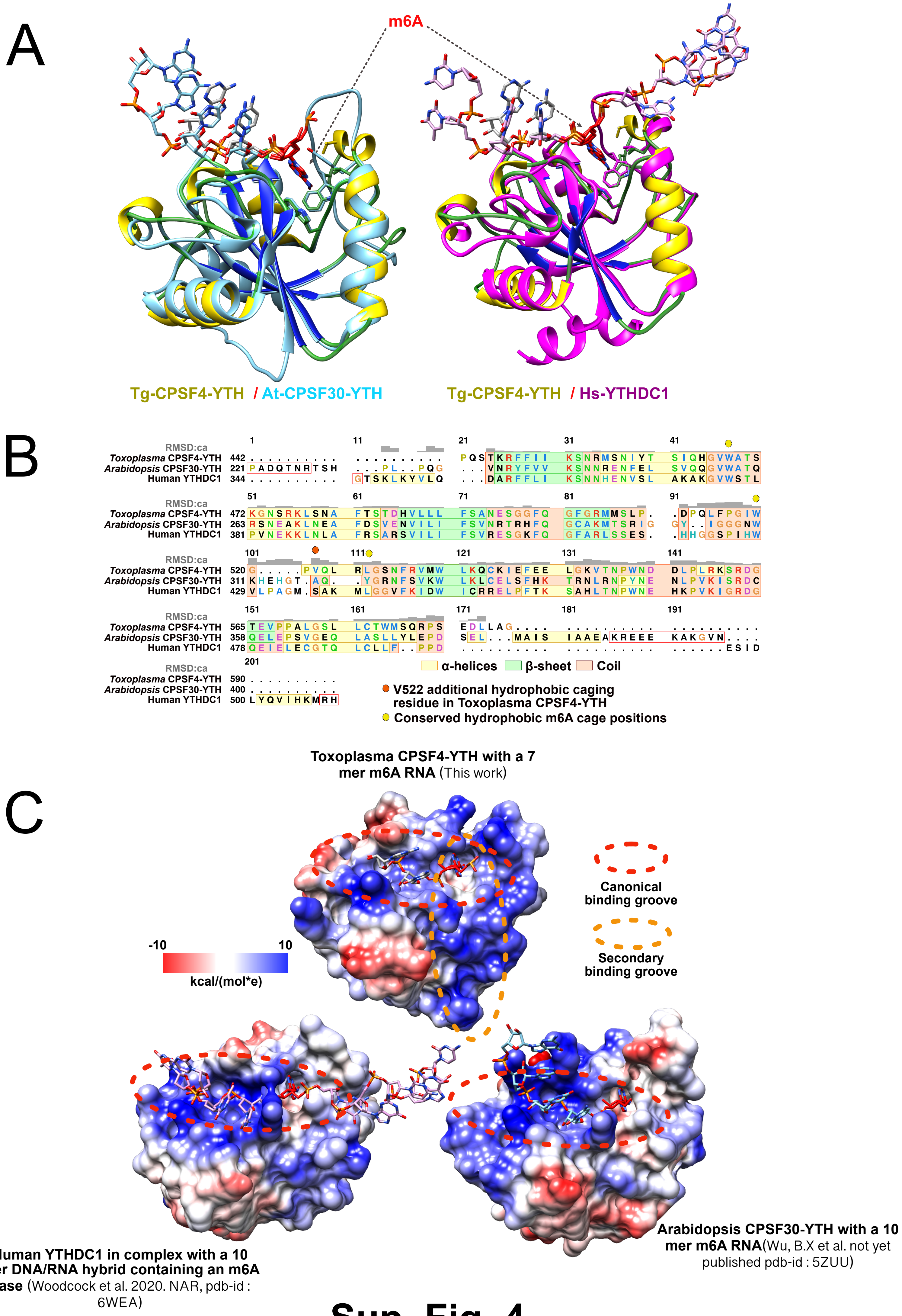
A
m6A
Tg-CPSF4-YTH / At-CPSF30-YTH
Tg-CPSF4-YTH / Hs-YTHDC1
B
RMSD:ca
1 11 21 31 41
Toxoplasma CPSF4-YTH 442 .......... .......... PQSTKRFFII KSNRMSNIYT SIQHGVWATS
Arabidopsis CPSF30-YTH 221 PADQTNRTSH ...PL..PQG ...VNRYFVV KSNNRENFEL SVQQGVWATQ
Human YTHDC1 344 .......... GTSKLKYVLQ ...DARFFLI KSNNHENVSL AKAKGVWSTL
RMSD:ca
51 61 71 81 91
Toxoplasma CPSF4-YTH 472 KGNSRKLSNA FTSTDHVLLL FSANESGGFQ GFGRMMSLP. .DPQLFPGIW
Arabidopsis CPSF30-YTH 263 RSNEAKLNEA FDSVENVILI FSVNRTRHFQ GCAKMTSRIG GY..IGGGNW
Human YTHDC1 381 PVNEKKLNLA FRSARSVILI FSVRESGKFQ GFARLSSES. .HHGGSPIHW
RMSD:ca
101 111 121 131 141
Toxoplasma CPSF4-YTH 520 G.....PVQL RLGSNFRVMW LKQCKIEFEE LGKVTNPWND DLPLRKSRDG
Arabidopsis CPSF30-YTH 311 KHEHGT.AQ. .YGRNFSVKW LKLCELSFHK TRNLRNPYNE NLPVKISRDC
Human YTHDC1 429 VLPAGM.SAK MLGGVFKIDW ICRRELPFTK SAHLTNPWNE HKPVKIGRDG
RMSD:ca
151 161 171 181 191
Toxoplasma CPSF4-YTH 565 TEVPPALGSL LCTWMSQRPS EDLLAG.... .......... ..........
Arabidopsis CPSF30-YTH 358 QELEPSVGEQ LASLLYLEPD SEL...MAIS IAAEAKREEE KAKGVN....
Human YTHDC1 478 QEIELECGTQ LCLLF..PPD .......... .......... ......ESID
RMSD:ca
201
Toxoplasma CPSF4-YTH 590 ..........
Arabidopsis CPSF30-YTH 400 ..........
Human YTHDC1 500 LYQVIHKMRH
α-helices β-sheet Coil
V522 additional hydrophobic caging residue in Toxoplasma CPSF4-YTH
Conserved hydrophobic m6A cage positions
C
Toxoplasma CPSF4-YTH with a 7 mer m6A RNA (This work)
Canonical binding groove
Secondary binding groove
-10
10
kcal/(mol*e)
Human YTHDC1 in complex with a 10 mer DNA/RNA hybrid containing an m6A base (Woodcock et al. 2020. NAR, pdb-id : 6WEA)
Arabidopsis CPSF30-YTH with a 10 mer m6A RNA(Wu, B.X et al. not yet published pdb-id : 5ZUU)

**Sup. Fig. 4**

A

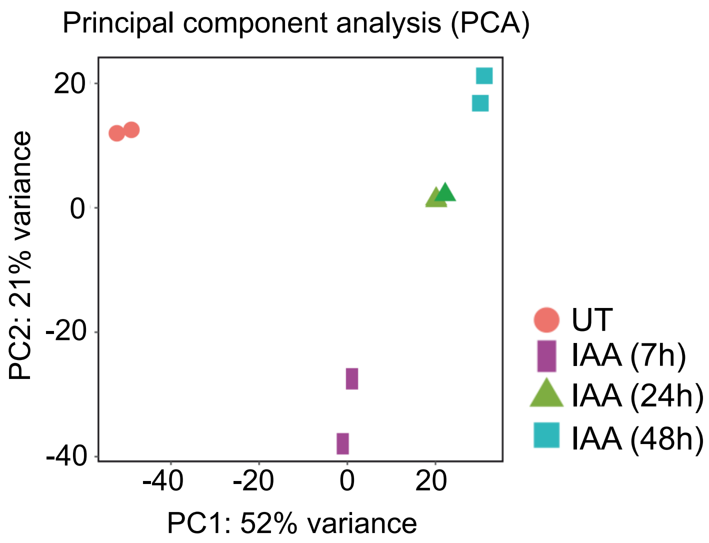
Principal component analysis (PCA)
20
0
-20
-40
-40
-20
0
20
PC2: 21% variance
PC1: 52% variance
UT
IAA (7h)
IAA (24h)
IAA (48h)

B

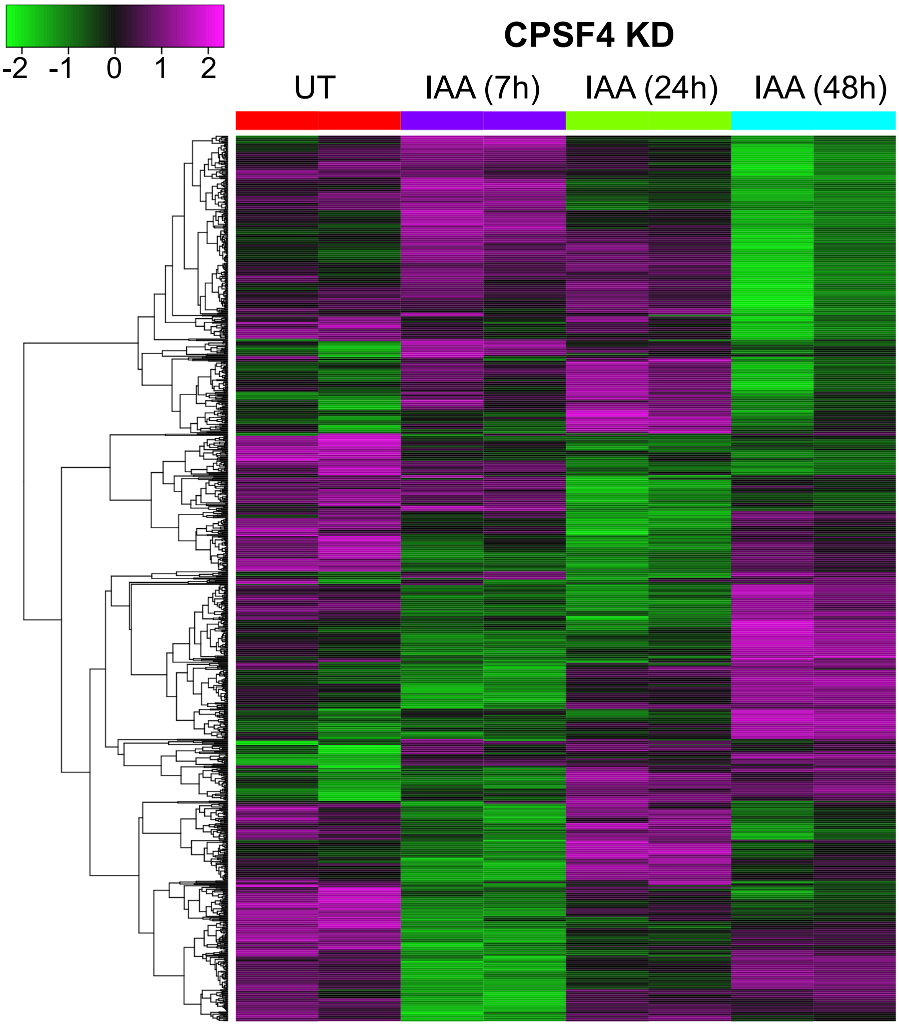
-2
-1
0
1
2
CPSF4 KD
UT
IAA (7h)
IAA (24h)
IAA (48h)

**Sup. Fig. 5**

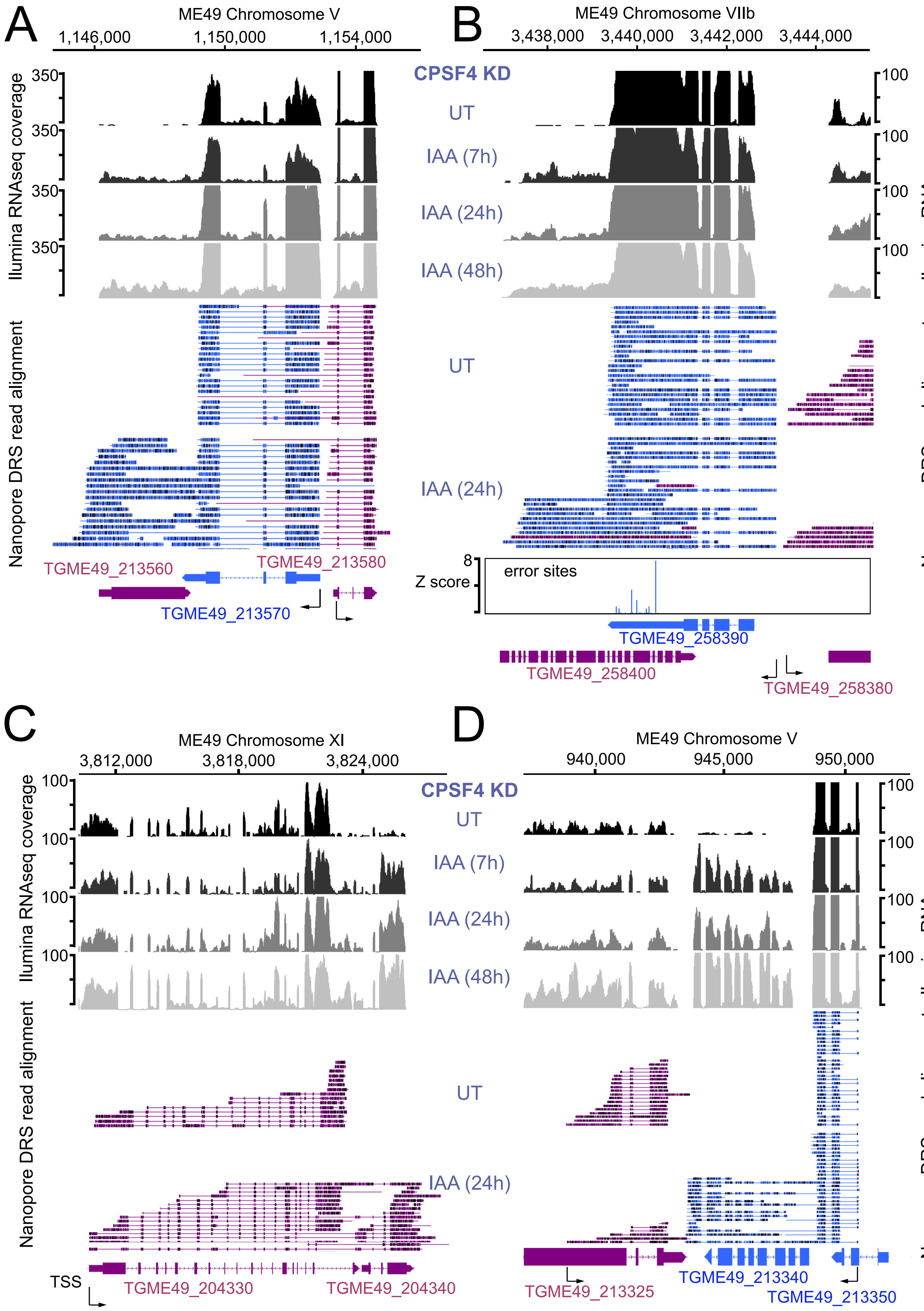

A
ME49 Chromosome V
1,146,000
1,150,000
1,154,000
Ilumina RNAseq coverage
350
Nanopore DRS read alignment
TGME49_213560
TGME49_213580
TGME49_213570
B
ME49 Chromosome VIIb
3,438,000
3,440,000
3,442,000
3,444,000
CPSF4 KD
UT
IAA (7h)
IAA (24h)
IAA (48h)
100
Ilumina RNAseq coverage
Nanopore DRS read alignment
8
Z score
error sites
TGME49_258390
TGME49_258400
TGME49_258380
C
ME49 Chromosome XI
3,812,000
3,818,000
3,824,000
100
Ilumina RNAseq coverage
Nanopore DRS read alignment
TSS
TGME49_204330
TGME49_204340
D
ME49 Chromosome V
940,000
945,000
950,000
CPSF4 KD
UT
IAA (7h)
IAA (24h)
IAA (48h)
100
Illumina RNAseq coverage
Nanopore DRS read alignment
TGME49_213325
TGME49_213340
TGME49_213350


**Sup. Fig. 6**

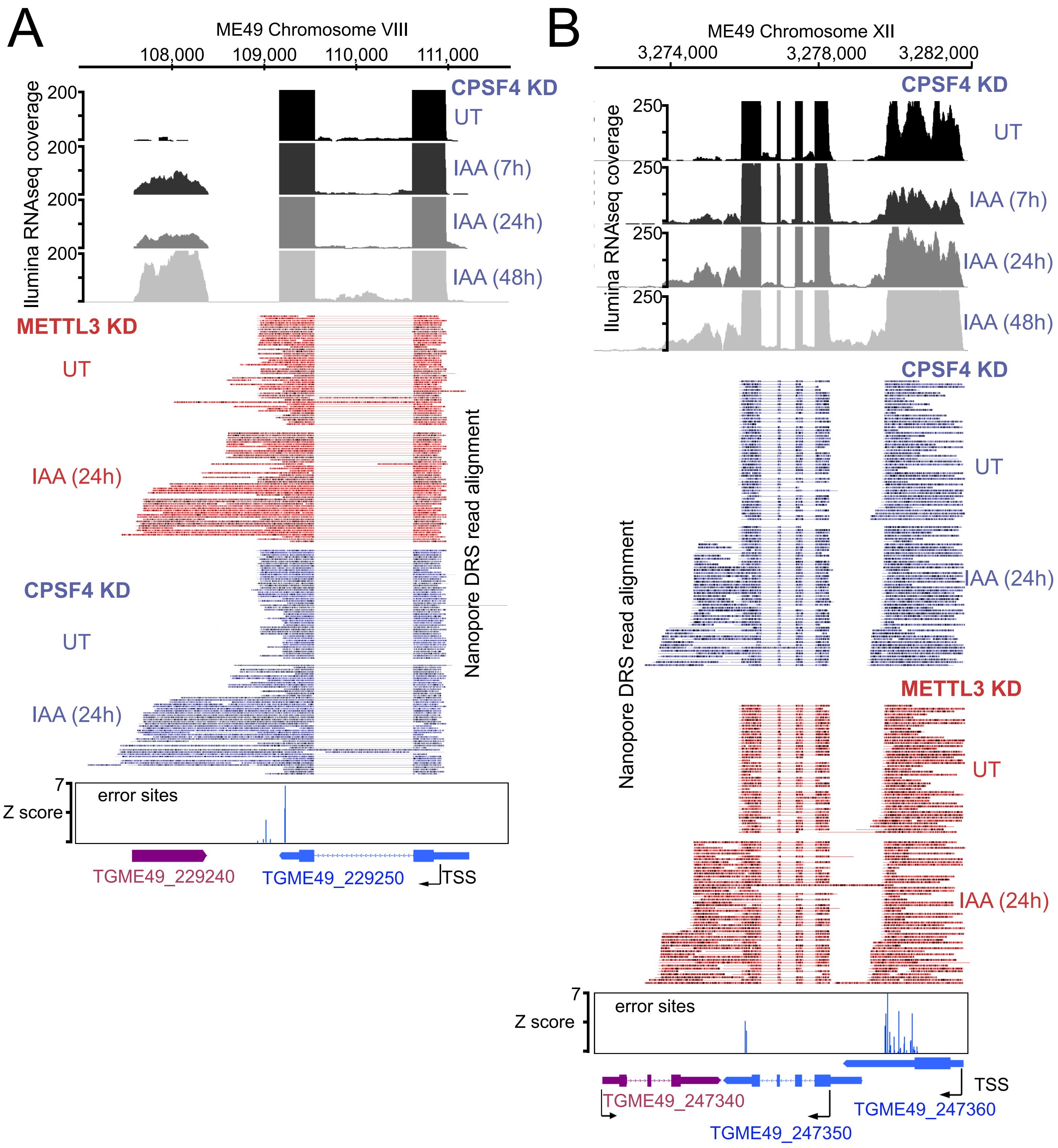
A
ME49 Chromosome VIII
108,000
109,000
110,000
111,000
Ilumina RNAseq coverage
200
CPSF4 KD
UT
IAA (7h)
IAA (24h)
IAA (48h)
METTL3 KD
UT
IAA (24h)
CPSF4 KD
UT
IAA (24h)
Nanopore DRS read alignment
7
Z score
error sites
TGME49_229240
TGME49_229250
TSS
B
ME49 Chromosome XII
3,274,000
3,278,000
3,282,000
CPSF4 KD
250
Ilumina RNAseq coverage
UT
IAA (7h)
IAA (24h)
IAA (48h)
CPSF4 KD
UT
IAA (24h)
Nanopore DRS read alignment
METTL3 KD
UT
IAA (24h)
7
Z score
error sites
TSS
TGME49_247340
TGME49_247360
TGME49_247350

### Sup. Fig. 7

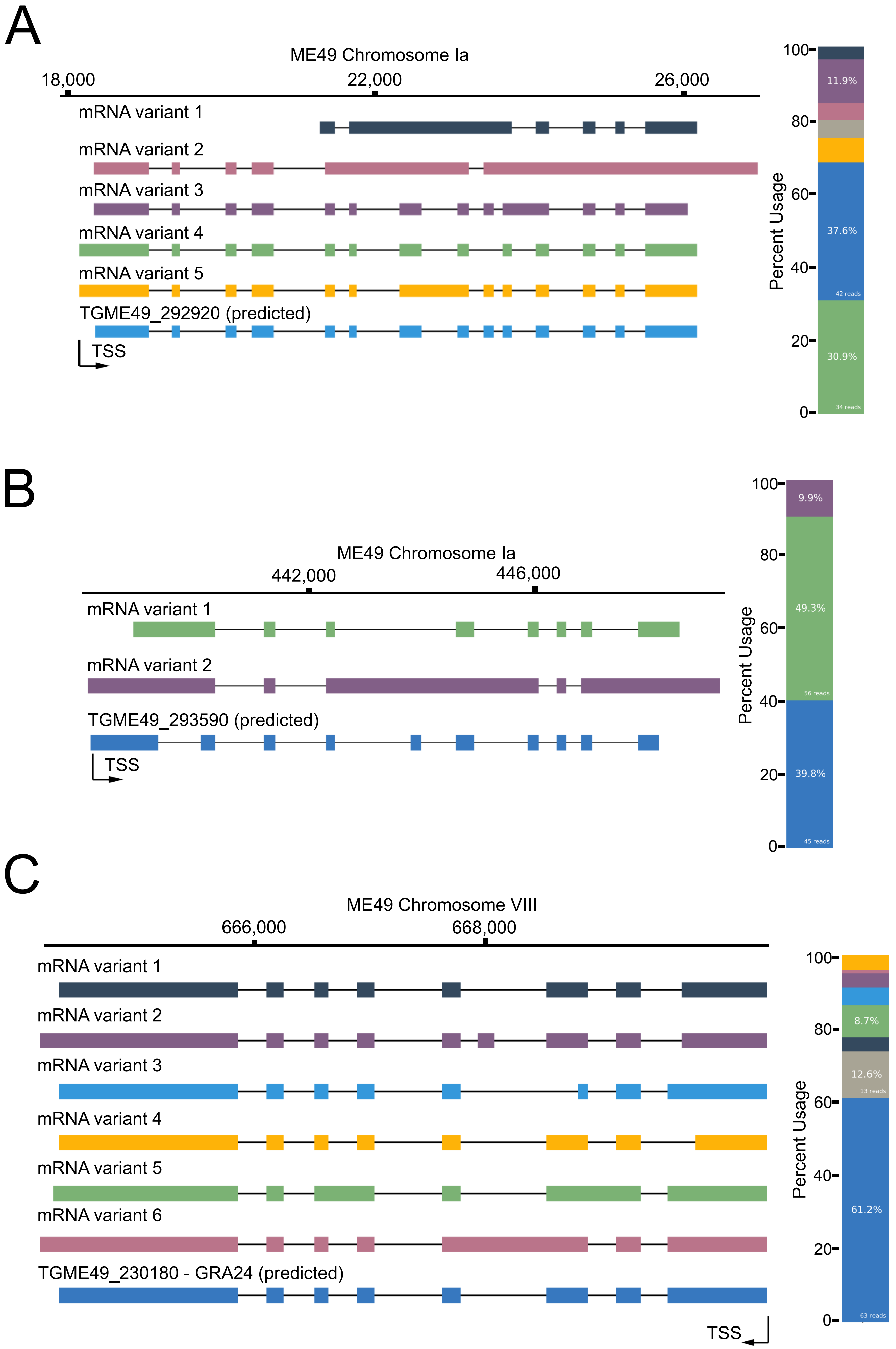
A
ME49 Chromosome Ia
18,000
22,000
26,000
mRNA variant 1
mRNA variant 2
mRNA variant 3
mRNA variant 4
mRNA variant 5
TGME49_292920 (predicted)
TSS
Percent Usage
100
80
60
40
20
0
11.9%
37.6%
42 reads
30.9%
34 reads
B
ME49 Chromosome Ia
442,000
446,000
mRNA variant 1
mRNA variant 2
TGME49_293590 (predicted)
TSS
Percent Usage
100
80
60
40
20
0
9.9%
49.3%
56 reads
39.8%
45 reads
C
ME49 Chromosome VIII
666,000
668,000
mRNA variant 1
mRNA variant 2
mRNA variant 3
mRNA variant 4
mRNA variant 5
mRNA variant 6
TGME49_230180 - GRA24 (predicted)
TSS
Percent Usage
100
80
60
40
20
0
8.7%
12.6%
13 reads
61.2%
63 reads

### Sup. Fig. 8

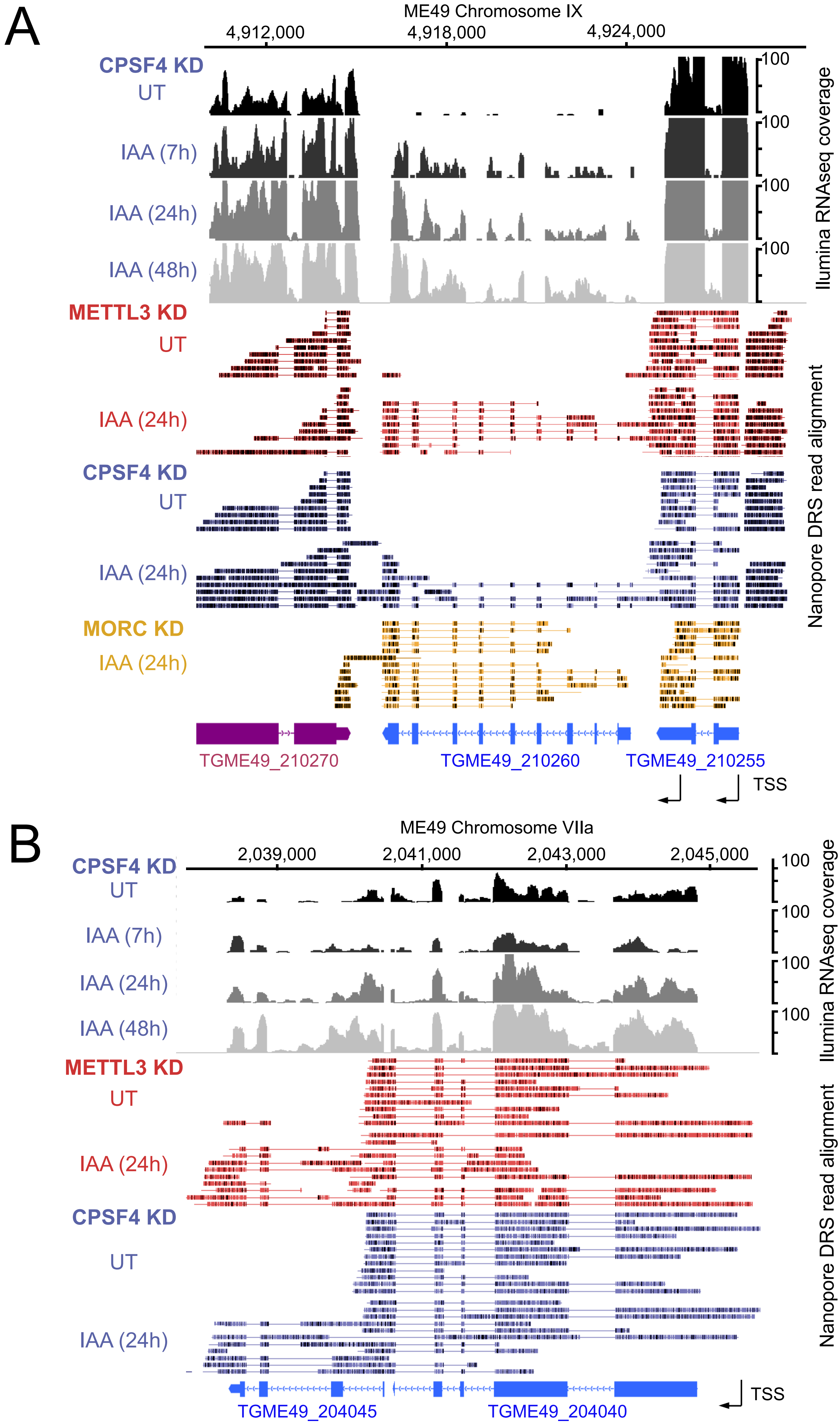
A
ME49 Chromosome IX
4,912,000
4,918,000
4,924,000
CPSF4 KD
UT
IAA (7h)
IAA (24h)
IAA (48h)
100
Ilumina RNAseq coverage
METTL3 KD
UT
IAA (24h)
CPSF4 KD
UT
IAA (24h)
MORC KD
IAA (24h)
Nanopore DRS read alignment
TGME49_210270
TGME49_210260
TGME49_210255
TSS
B
ME49 Chromosome VIIa
2,039,000
2,041,000
2,043,000
2,045,000
CPSF4 KD
UT
IAA (7h)
IAA (24h)
IAA (48h)
100
Ilumina RNAseq coverage
METTL3 KD
UT
IAA (24h)
CPSF4 KD
UT
IAA (24h)
Nanopore DRS read alignment
TGME49_204045
TGME49_204040
TSS

**Sup. Fig. 9**

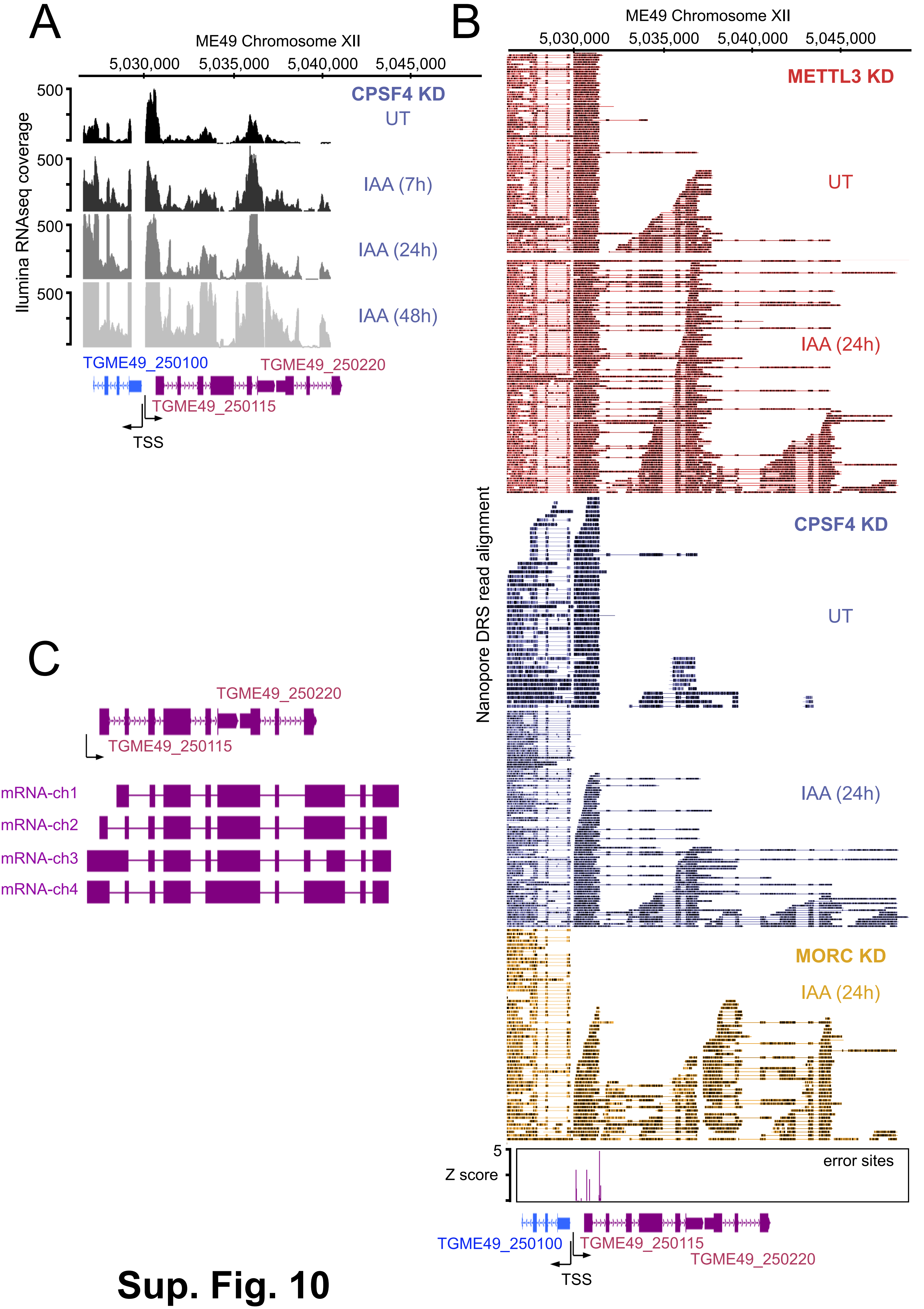
A
ME49 Chromosome XII
5,030,000
5,035,000
5,040,000
5,045,000
Ilumina RNAseq coverage
500
CPSF4 KD
UT
IAA (7h)
IAA (24h)
IAA (48h)
TGME49_250100
TGME49_250220
TGME49_250115
TSS
B
ME49 Chromosome XII
METTL3 KD
UT
IAA (24h)
Nanopore DRS read alignment
CPSF4 KD
UT
IAA (24h)
MORC KD
IAA (24h)
5
Z score
error sites
TGME49_250100
TGME49_250115
TGME49_250220
TSS
C
TGME49_250220
TGME49_250115
mRNA-ch1
mRNA-ch2
mRNA-ch3
mRNA-ch4

### Sup. Fig. 10

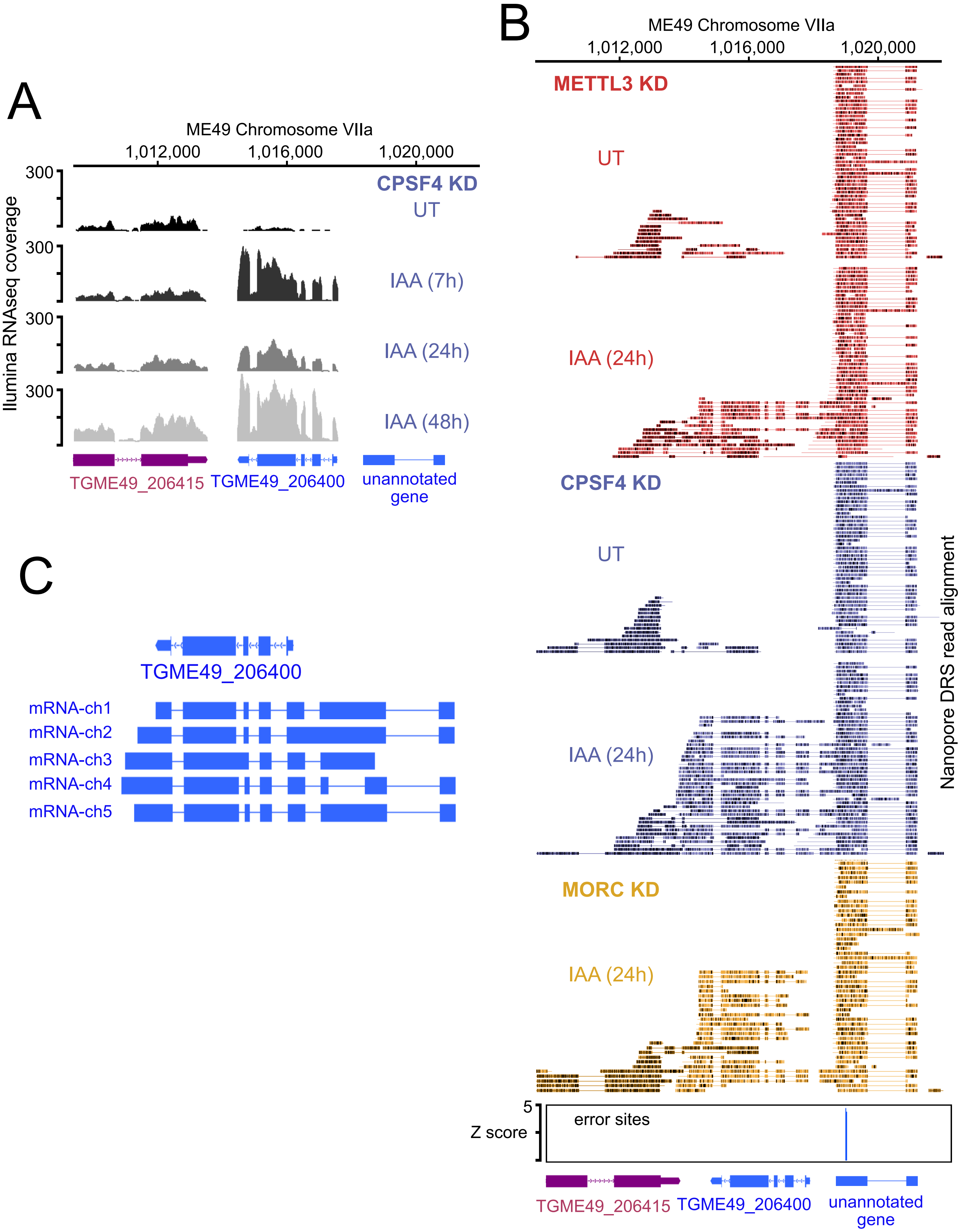
A
ME49 Chromosome VIIa
1,012,000
1,016,000
1,020,000
Ilumina RNAseq coverage
300
CPSF4 KD
UT
IAA (7h)
IAA (24h)
IAA (48h)
TGME49_206415
TGME49_206400
unannotated gene
B
ME49 Chromosome VIIa
1,012,000
1,016,000
1,020,000
METTL3 KD
UT
IAA (24h)
CPSF4 KD
UT
IAA (24h)
MORC KD
IAA (24h)
Nanopore DRS read alignment
5
Z score
error sites
TGME49_206415
TGME49_206400
unannotated gene
C
TGME49_206400
mRNA-ch1
mRNA-ch2
mRNA-ch3
mRNA-ch4
mRNA-ch5

**Sup. Fig. 11**

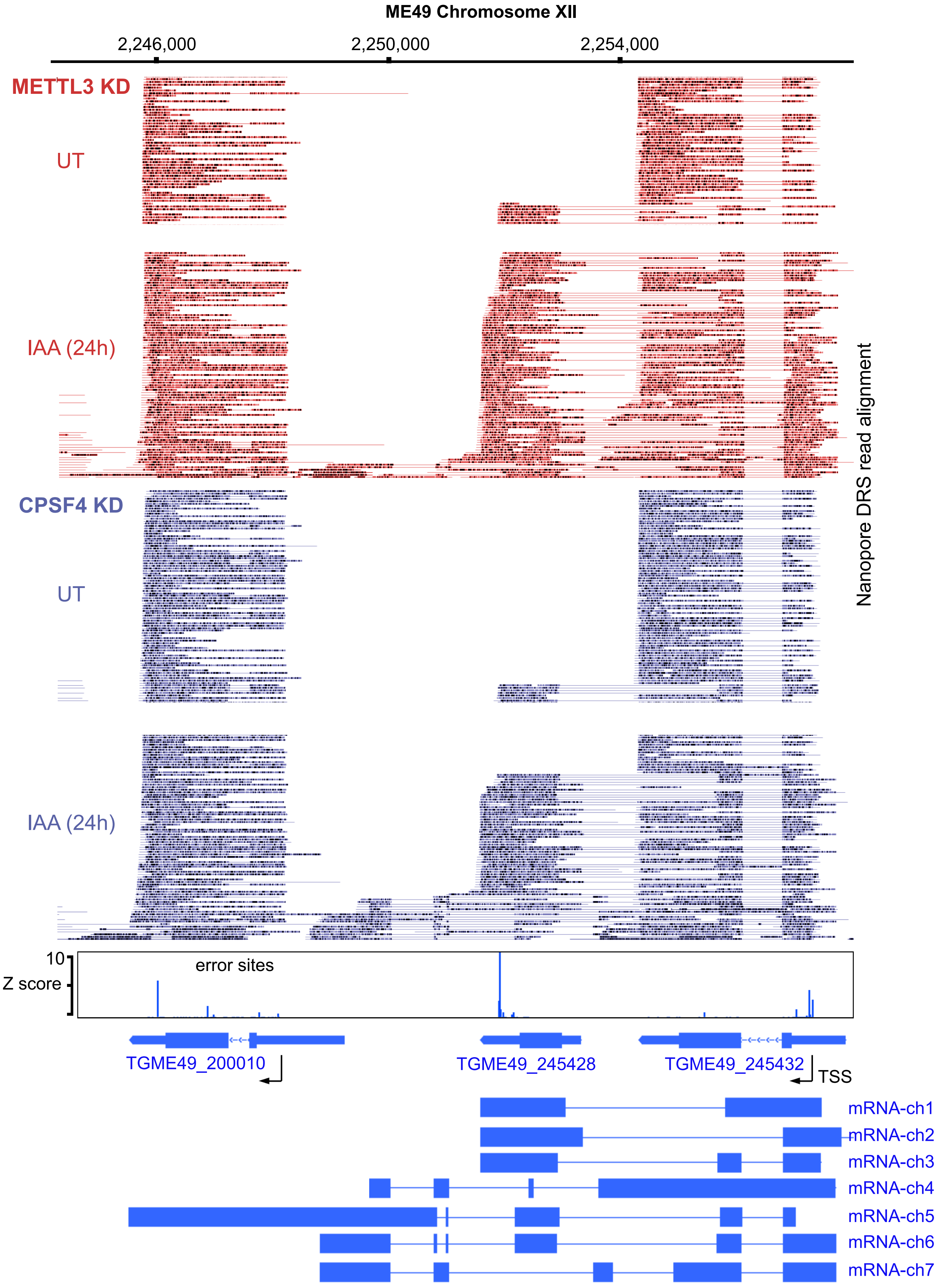
ME49 Chromosome XII
2,246,000
2,250,000
2,254,000
METTL3 KD
UT
IAA (24h)
CPSF4 KD
UT
IAA (24h)
Nanopore DRS read alignment
10
Z score
error sites
TGME49_200010
TGME49_245428
TGME49_245432
TSS
mRNA-ch1
mRNA-ch2
mRNA-ch3
mRNA-ch4
mRNA-ch5
mRNA-ch6
mRNA-ch7

### Sup. Fig. 12

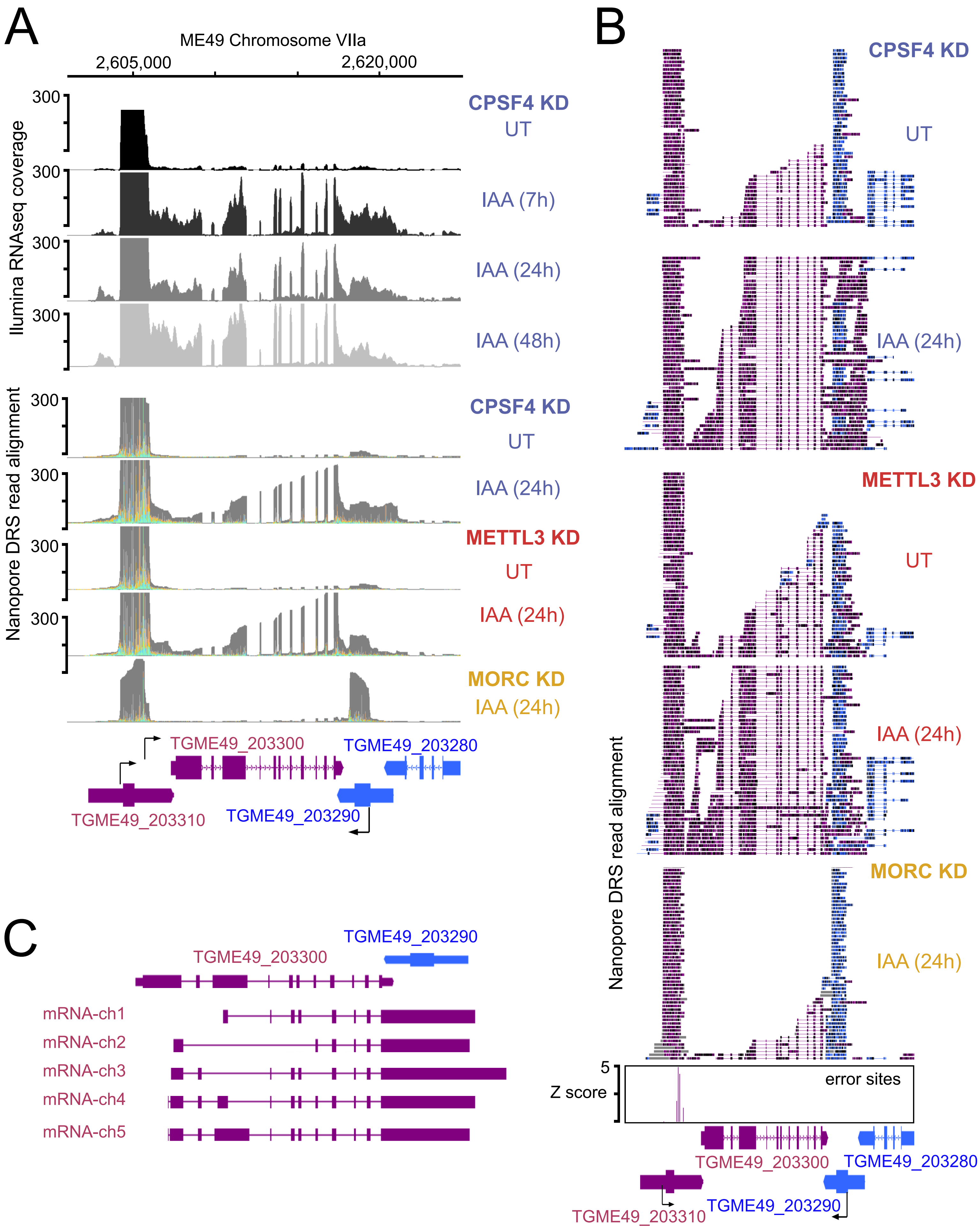
A
ME49 Chromosome VIIa
2,605,000
2,620,000
Illumina RNAseq coverage
300
CPSF4 KD
UT
IAA (7h)
IAA (24h)
IAA (48h)
Nanopore DRS read alignment
CPSF4 KD
UT
IAA (24h)
METTL3 KD
UT
IAA (24h)
MORC KD
IAA (24h)
TGME49_203300
TGME49_203280
TGME49_203310
TGME49_203290
B
CPSF4 KD
UT
IAA (24h)
METTL3 KD
UT
IAA (24h)
MORC KD
IAA (24h)
Nanopore DRS read alignment
5
Z score
error sites
TGME49_203300
TGME49_203280
TGME49_203310
TGME49_203290
C
TGME49_203290
TGME49_203300
mRNA-ch1
mRNA-ch2
mRNA-ch3
mRNA-ch4
mRNA-ch5

**Sup. Fig. 13**

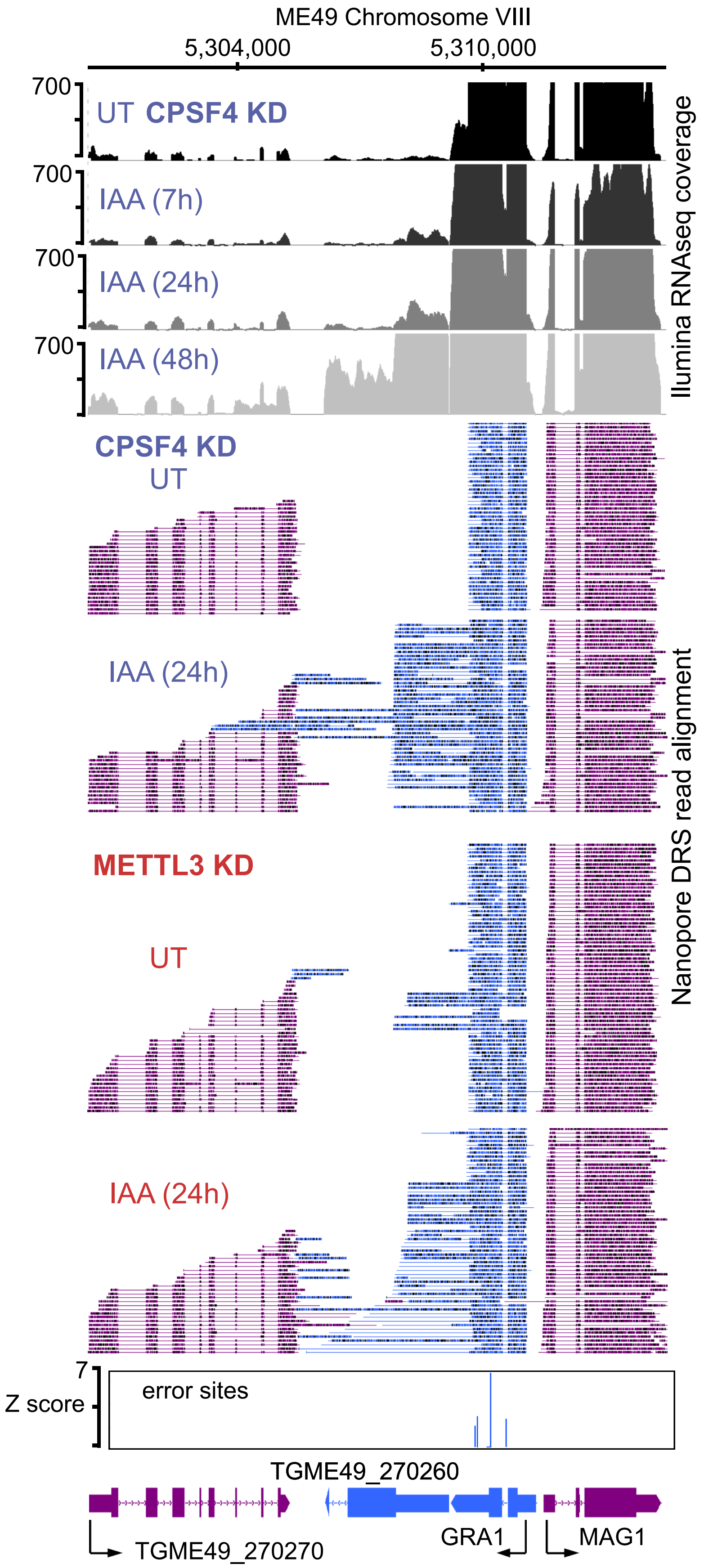
ME49 Chromosome VIII
5,304,000
5,310,000
700
UT CPSF4 KD
700
IAA (7h)
700
IAA (24h)
700
IAA (48h)
Ilumina RNAseq coverage
CPSF4 KD
UT
IAA (24h)
METTL3 KD
UT
IAA (24h)
Nanopore DRS read alignment
7
Z score
error sites
TGME49_270260
TGME49_270270
GRA1
MAG1

### Sup. Fig. 14

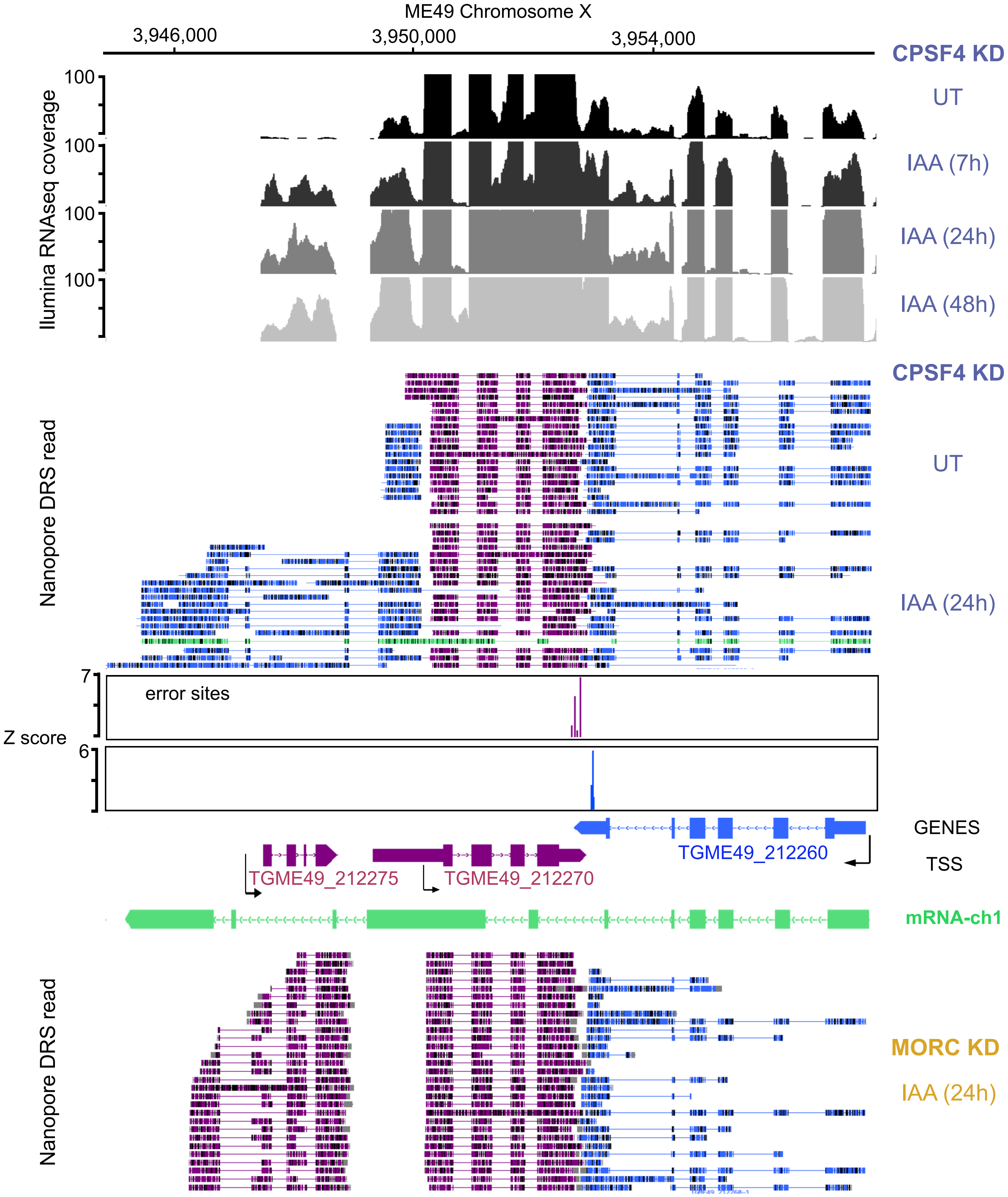

ME49 Chromosome X
3,946,000
3,950,000
3,954,000
CPSF4 KD
UT
IAA (7h)
IAA (24h)
IAA (48h)
100
100
100
100
Ilumina RNAseq coverage
CPSF4 KD
UT
IAA (24h)
Nanopore DRS read
error sites
Z score
7
6
GENES
TSS
TGME49_212260
TGME49_212275
TGME49_212270
mRNA-ch1
Nanopore DRS read
MORC KD
IAA (24h)


### Sup. Fig. 15

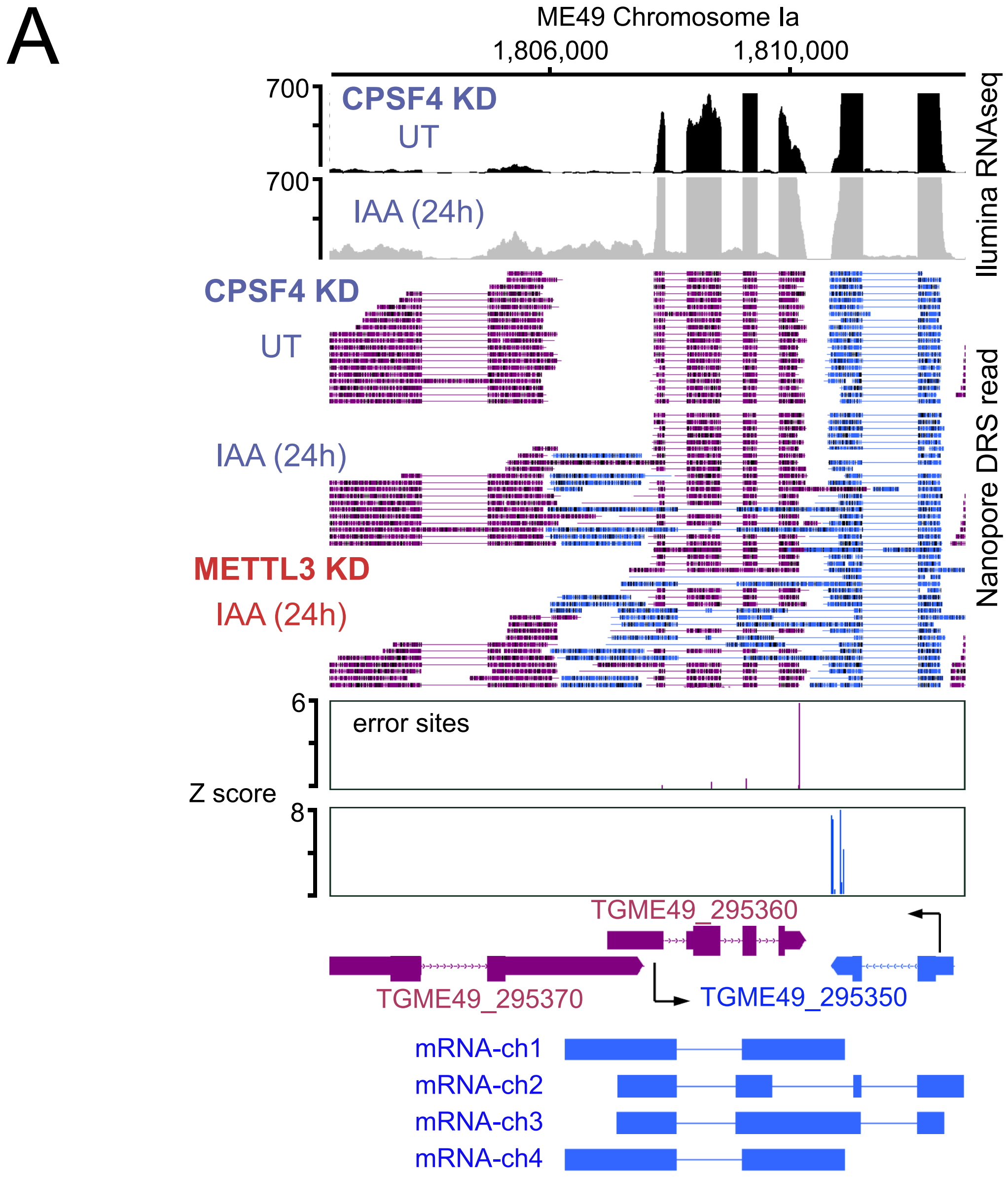
A
ME49 Chromosome Ia
1,806,000
1,810,000
700
CPSF4 KD
UT
700
IAA (24h)
Ilumina RNAseq
CPSF4 KD
UT
IAA (24h)
METTL3 KD
IAA (24h)
Nanopore DRS read
6
error sites
Z score
8
TGME49_295360
TGME49_295370
TGME49_295350
mRNA-ch1
mRNA-ch2
mRNA-ch3
mRNA-ch4

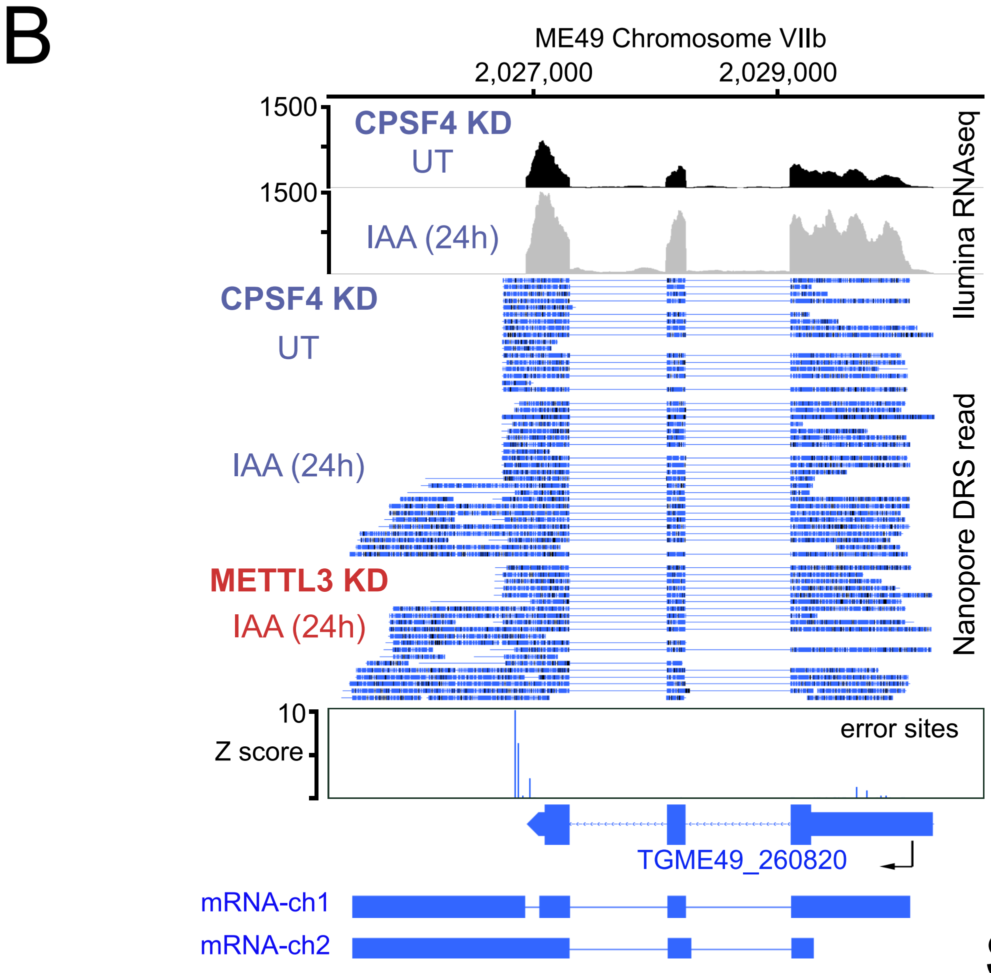
B
ME49 Chromosome VIIb
2,027,000
2,029,000
1500
CPSF4 KD
UT
1500
IAA (24h)
Ilumina RNAseq
CPSF4 KD
UT
IAA (24h)
METTL3 KD
IAA (24h)
Nanopore DRS read
10
Z score
error sites
TGME49_260820
mRNA-ch1
mRNA-ch2

Sup. Fig. 16

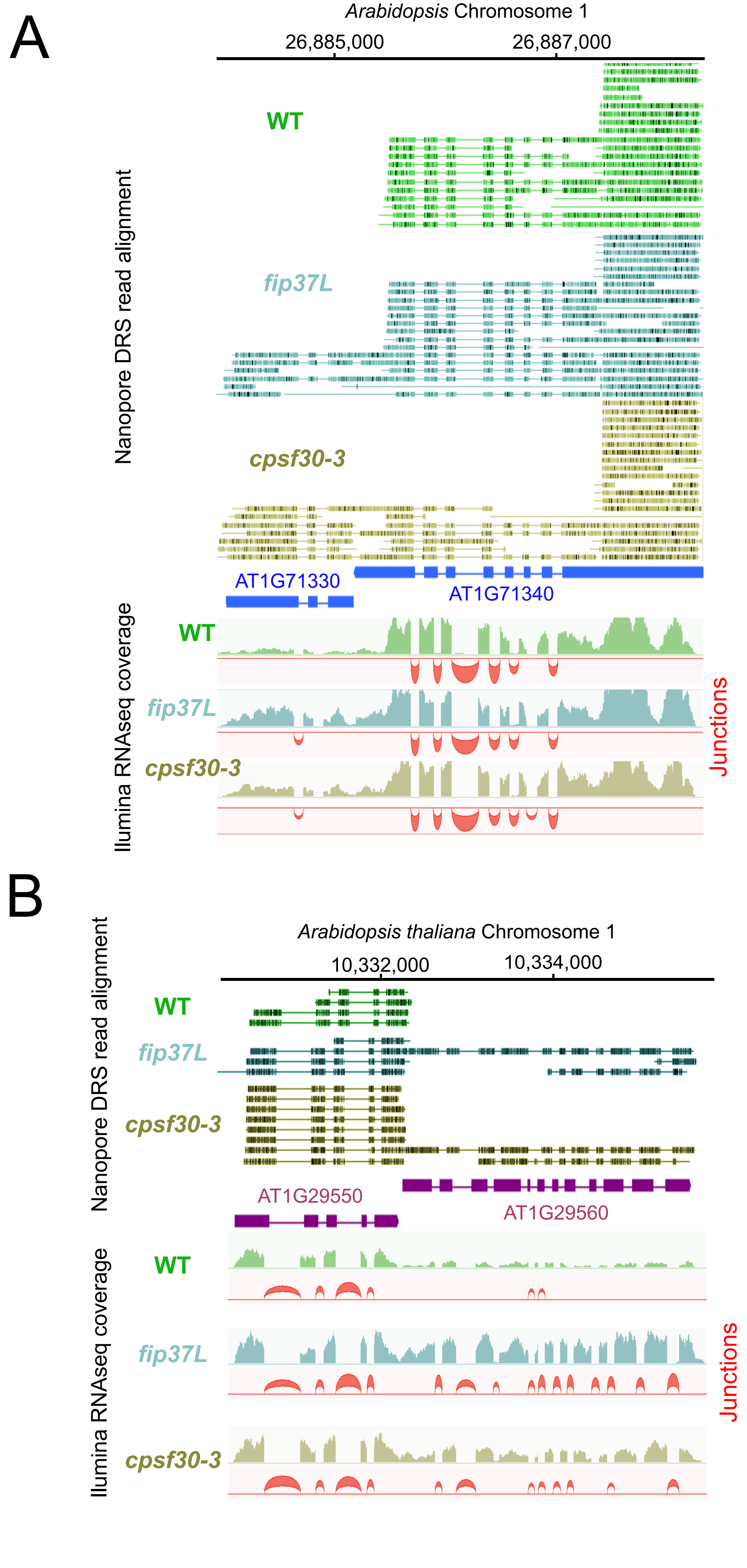
A
Arabidopsis Chromosome 1
26,885,000
26,887,000
WT
fip37L
cpsf30-3
Nanopore DRS read alignment
AT1G71330
AT1G71340
WT
fip37L
cpsf30-3
Ilumina RNAseq coverage
Junctions
B
Arabidopsis thaliana Chromosome 1
10,332,000
10,334,000
WT
fip37L
cpsf30-3
Nanopore DRS read alignment
AT1G29550
AT1G29560
WT
fip37L
cpsf30-3
Ilumina RNAseq coverage
Junctions

**Sup. Fig. 17**

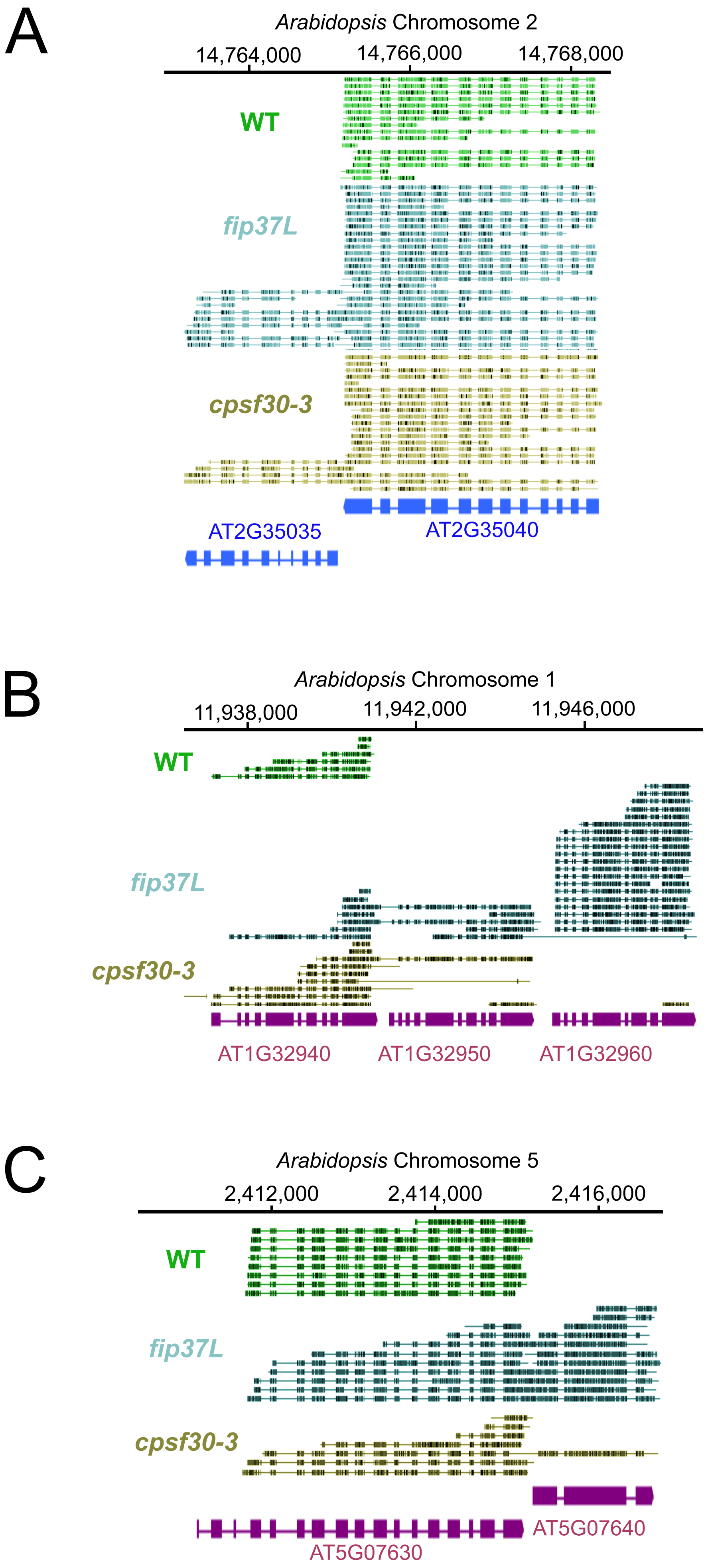
A
Arabidopsis Chromosome 2
14,764,000
14,766,000
14,768,000
WT
fip37L
cpsf30-3
AT2G35035
AT2G35040
B
Arabidopsis Chromosome 1
11,938,000
11,942,000
11,946,000
WT
fip37L
cpsf30-3
AT1G32940
AT1G32950
AT1G32960
C
Arabidopsis Chromosome 5
2,412,000
2,414,000
2,416,000
WT
fip37L
cpsf30-3
AT5G07640
AT5G07630

**Sup. Fig. 18**
